# Monthly physical activity and storage time affect enzyme acetylcholinesterase activity in Plasma and Whole blood samples of healthy elderly subjects

**DOI:** 10.1101/2020.09.16.299610

**Authors:** Snežana M. Jovičić

**Affiliations:** Department of Genetics, Faculty of Biology, University of Belgrade, Studentski trg 16, 11000 Belgrade, Serbia

**Keywords:** physical activity, pre-analytical factors, enzyme acetylcholinesterase, inhibitor Eserin and BW284c51, healthy elderly individuals

## Abstract

**Objective:** Enzyme Acetylcholinesterase is a potential marker for ageing, disease onset, progression and therapy. Physical activity and storage time affect the variable range, quality data prevalence and variability. The study aims to examine the effect of one and two months of storage time on protein concentration, acetylcholinesterase activity and inhibitor efficacy in subjects subjected to 3 months of physical activity.

**Method:** The value of AChE activity was measured spectrophotometrically by Elman et al. (1961). The obtained results were processed by the statistical program SPSS v 23.0.

**Results:** Group of 32 healthy subjects showed a significant difference (P<0.01) for protein concentration and acetylcholinesterase activity in Plasma, inhibitor Eserine efficacy in Plasma and Whole blood before and after physical activity. Based on Age, a significant difference is present for Height (P<0.01), Muscle mass (P<0.01) and protein concentration (P<0.05) in Whole blood after physical activity and storage time. A statistically significant difference between Gender exists for Age (P<0.01), Height (P<0.01), Weight (P<0.05), Muscle mass (P<0.05), protein concentration (P<0.05) in whole blood and acetylcholinesterase activity (P<0.05) in Plasma after physical activity.

**Conclusions:** Preliminary and descriptive data indicate no difference in acetylcholinesterase activity between Age groups. Physical activity and storage time influence the change of protein concentration, enzyme activity and inhibitor Eserine efficacy in healthy elderly subjects. Future studies should compare disease and healthy participants.

## Introduction

Organism health defines as complete physical, mental and social well-being measured by the absence of disease and infirmity [World Health Organization, 2021]. Life expectancy assesses the health of a population-based on Gender and Age [Parker M, 2020]. Factors that influence health are in and beyond the power of the individual. Biological and non-biological factors affect the values of the examined biomarkers in populations [Freitas Leal JK, 2017]. The most important are genetics, public policies and regulations (vaccination campaigns), health (financing, diagnosis and treatment of diseases), life habits (smoking, physical activity, hydration of the organism), social and environmental factors (bioterrorism, hygiene, environmental pollution, infections, drugs, nutrition, religion and ethnicity) [Radovanovic-Nenadic U, 2020; Ramaswami R, 2018].

Ageing is a universal process in all living creatures and induces a change in organism physiological function under diverse conditions [Cinar D, 2015]. Ageing is linked to various disease onsets and progression in humans like cancer. Population study estimated 3.91 million new cases and 1.93 million deaths from cancer in Europe in 2018 [Ferlay J, 2018]. The most frequently seen cancer in the elderly population is lung, breast, cervical, colorectal and prostate cancer [Cinar D, 2015]. Cholinergic transmission disorder is a well-established pathological pattern of diseases [Chang EH, 2019]. Cholinesterases are neurotransmitters that catalyze the hydrolysis of the cholinergic neurotransmitter acetylcholine [Lockridge O, 2016]. Enzyme acetylcholinesterase (AChE) applies in clinical practice for disease diagnosis, therapy and treatment [Lockridge O, 2016]. Treatment is poorly effective or ineffective in most cases [Chang EH, 2019]. Treatment reduces symptoms and does not cure the disease [Chang EH, 2019]. Physical activity applies as a therapy for chronic diseases [Lockridge O, 2016].

Knowledge of all homeostatic conditions in the healthy human body and pre-analytical variables influences data interpretation quality [Islam MS, 2018]. Pre-analytical factors like sample sorting, recording, preservation, storage time, material, method, environmental factors (hydration, physical activity, food intake) enable gathering high-quality sample information [Islam MS, 2018]. Various studies have analyzed the influence of pre-analytical factors such as storage time and physical activity on biomarker range and repeatability of results. The beneficial and harmful effect of physical activity is studied [Joyner MJ, 2015; Simioni C, 2018]. During sample storage, erythrocyte age and enzyme AChE degradate [Freitas Leal JK, 2017]. Changes in AChE enzyme activity over several days, months, and years of storage are known in healthy subjects [Freitas Leal JK, 2017, Crane CR, 1970].

There is a lack of knowledge of the influence of pre-analytical factors, physical activity and time storage on enzyme AChE activity in healthy elderly subjects. These can be a further guiding stone for disease research in the elderly population. The current study hypothesizes **t**hat changes in the blood AChE activity are associated with physical activity and sample storage time in healthy elderly subjects. The objective was to test the hypothesis, examine and discuss the effect of monthly physical exercise and storage time on protein concentration, AChE activity and inhibitor Eserine and BW284c51 efficacy in healthy elderly subjects.

## Materials and Methods

### Study group and Human biological material

Participants differ based on Gender, Age and anthropometric measurements, protein concentration, enzyme AChE activity, inhibitor efficacy in whole blood (WB) and PL samples of healthy individuals. All procedures contributing to this work comply with the ethical standards of the relevant national and institutional committees on human experimentation and with the Helsinki Declaration of 1975, as revised in 2008. All procedures involving human subjects/patients were approved by the National Medical Ethics Committee of the Republic of Slovenia, Institute of Clinical Neurophysiology, number 82/07/14.

Male and female participants divide into two age groups, group A with the range of 60-70 years and group B with 70-80 years. Female participants were prevalent in both age groups (group A: 60%/81.25% and group B: 14%/18.75%). The total number at the beginning of the study consisted of 64 healthy elderly participants. Female participants (49, 76.56%) were numerous to males (15, 23.43%). The total mean Age for males and females was 68.625 years (min=59, max=85), SD=4.964. Male mean Age was 69.93 years (min=65, max=80), SD=4.69. Female mean Age was 68.22 years (min=59, max=85), SD=5.021. At the end of the study, 32 healthy elderly participants completed the survey process. Female participants (22, 68.75%) remain numerous to males (10, 31.25%). The total mean Age for males and females was 68.813 years (min=60, max=80), SD=4.41. Male mean Age was 70.8 years (min=65, max=80), SD=5.25. Female mean Age was 66.45 years (min=60, max=74), SD=3.27. Participant blood was sampled twice, at the beginning of the study and after 3. months of physical activity. Collected samples are stored in the refrigerator at −21°C for 2. months (1^st^ sampling) and 1. month (2^nd^ sampling) before analysis.

### A sampling of biological material

WB samples were collected from healthy individuals by phlebotomy using needles of 21 diameters (70 mm length, 0.4 mm inner diameter, Microlance, Becton Dickinson, Franklin Lakes, NJ, USA) and stored in four 2.7 mL vacutainers containing 270 µL trisodium citrate (0.109 mol/L) as an anticoagulant. From the subject biological material, PL is isolated.

### Fractioning of biological material

#### Plasma isolation

100µL of WB was sampled into the Eppendorf tube to measure cholinesterase activity. Blood was centrifuged at 1550 g, 37°C, for 20 minutes in a centric 200/R centrifuge (Domel doo, Železniki, Slovenia) to separate cells from PL by Thermo Fisher Scientific Protocol. PL sample fraction is removed and stored in a 2.7 mL vial without the presence of anticoagulants and analyzed without washing or further processing.

### Biochemical methodology

#### Measurement of protein concentration

To measure protein absorption on 96 well plates using Pierce BCA protein assay kit for one sample, triplicate reaction: sample reaction, negative control reaction, and standard curve (bovine serum albumin-BSA dilution of 2mg/ml, 1mg/ml, 0.5mg/ml, 0.25mg/ml, 0.125mg/ml, 0.0625 mg/ml) is performed. Depending on the type of reaction in each well, we had: 20 µL of sample/buffer/BSA dilution and 200 µL of a mixture of reagent A and reagent B in a ratio of 1:50). Samples incubated for 30 minutes at 37°C. The absorbance was measured at 550 nm using a spectrophotometer (BioTek, Cytation 3, Bad Friedrichshall, Germany) for one cycle (30 s).

### Measuring cholinesterase activity

In the selected samples (WB, PL), enzyme AChE activity was measured, following the Ellman method (Ellman et al. 1961) [Ellman GL, 1961]. The obtained samples dilute in potassium phosphate buffer (100 mM, pH=8.0). Into 96 well plates, three types of reactions (induced, inhibited and negative control reaction) are employed to measure cholinesterase activity. All the measurements performed at room temperature in triplicates. Depending on the reaction type in each well, we have different reagents. Induced/inhibited chemical reaction contains 100 µL of the diluted sample, 100 µL of Elman reagent (DTNB-5,5′-dithiobis (2-nitrobenzoic acid)) in 250mM potassium phosphate buffer, pH 7.4) without/with acetylthiocholine chloride substrate (1mM) and 20 µL of potassium phosphate buffer/inhibitor (potassium phosphate buffer/Eserine/BW284c51). The negative control reaction contains 120 µL potassium phosphate buffer and 100 µL Elman substrate reagents. Absorbance was measured at 420 nm using a spectrophotometer (BioTek, Cytation 3, Bad Friedrichshall, Germany) for 40 cycles (30 s interval per cycle, 20minutes).

### Statistical analysis

Data analysis was performed by IBM SPSS software version 23.0 for statistical analysis. Descriptive statistics, Shapiro-Wilk normality test, parametric and non-parametric statistical tests such as paired, independent and one sample T-test, Wilcoxon, Mann Whitney, Kruskal Walis is employed. Results display mean (SD) and p-value. A statistically significant correlation was assumed when P<0.05.

## Results

### Characteristics of the participant group

Descriptive statistics results in Table 1 shows the general characteristics of the participants. Participants differ based on Gender (Male and Female) and Age (Group A: 60-70 years; Group B: 70-80 years). Proposed values of protein concentration, enzyme AChE activity and inhibitor efficacy is subject to change. Participant protein concentration, enzyme AChE activity and inhibitor efficacy change before and after physical activity and storage time in healthy elderly participants are present in Figure 1-3. Shapiro-Wilk test indicated the presence of normality (P>0.05) for anthropometry measures (Height, Weight, Waist circumference), protein concentration (PL after and WB before and after physical activity) and enzyme AChE activity (PL after and WB before and after physical activity). Further parametric and non-parametric statistical tests apply. Logarithm transformation transforms non-parametric data to normal.

**Table 1.**
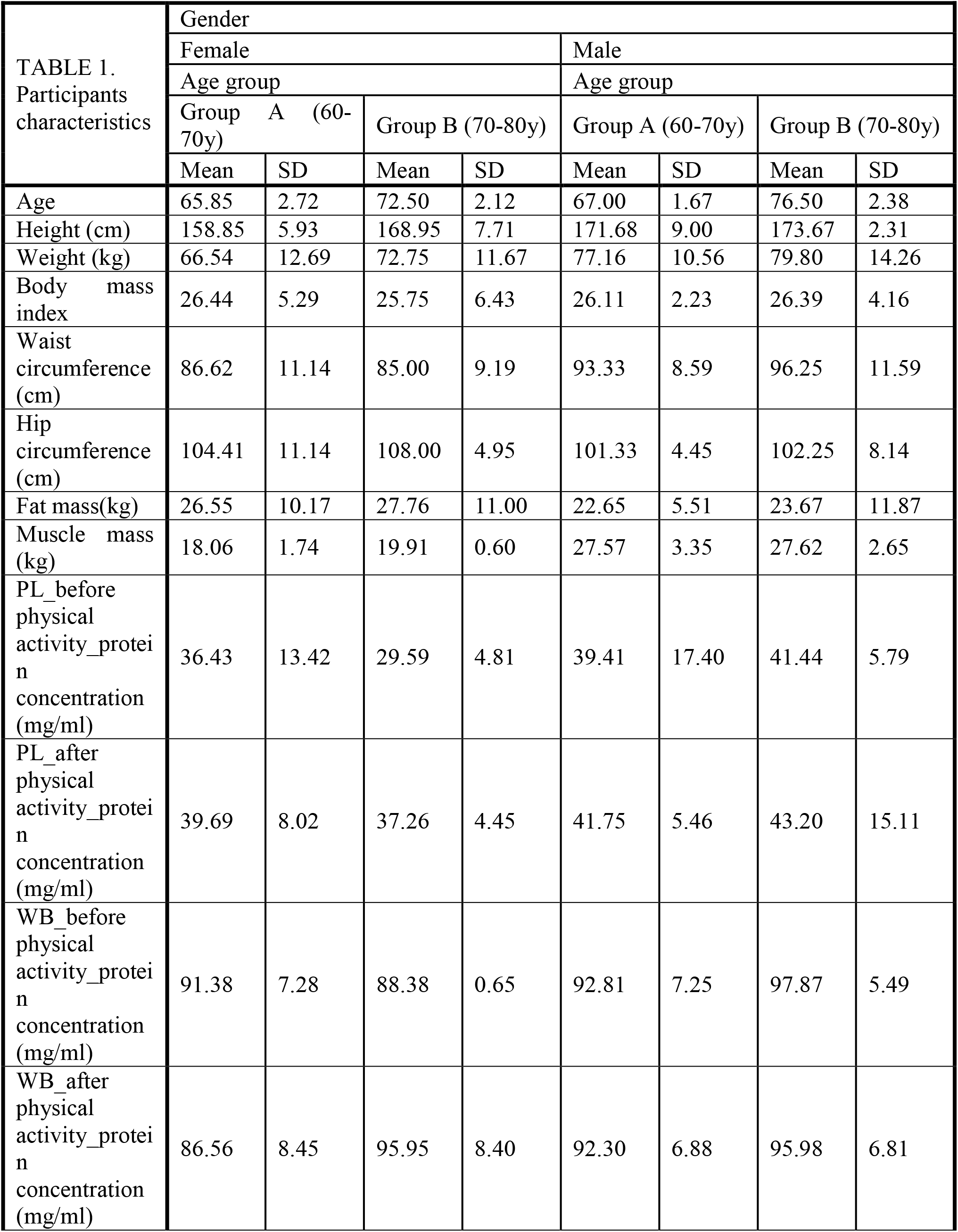
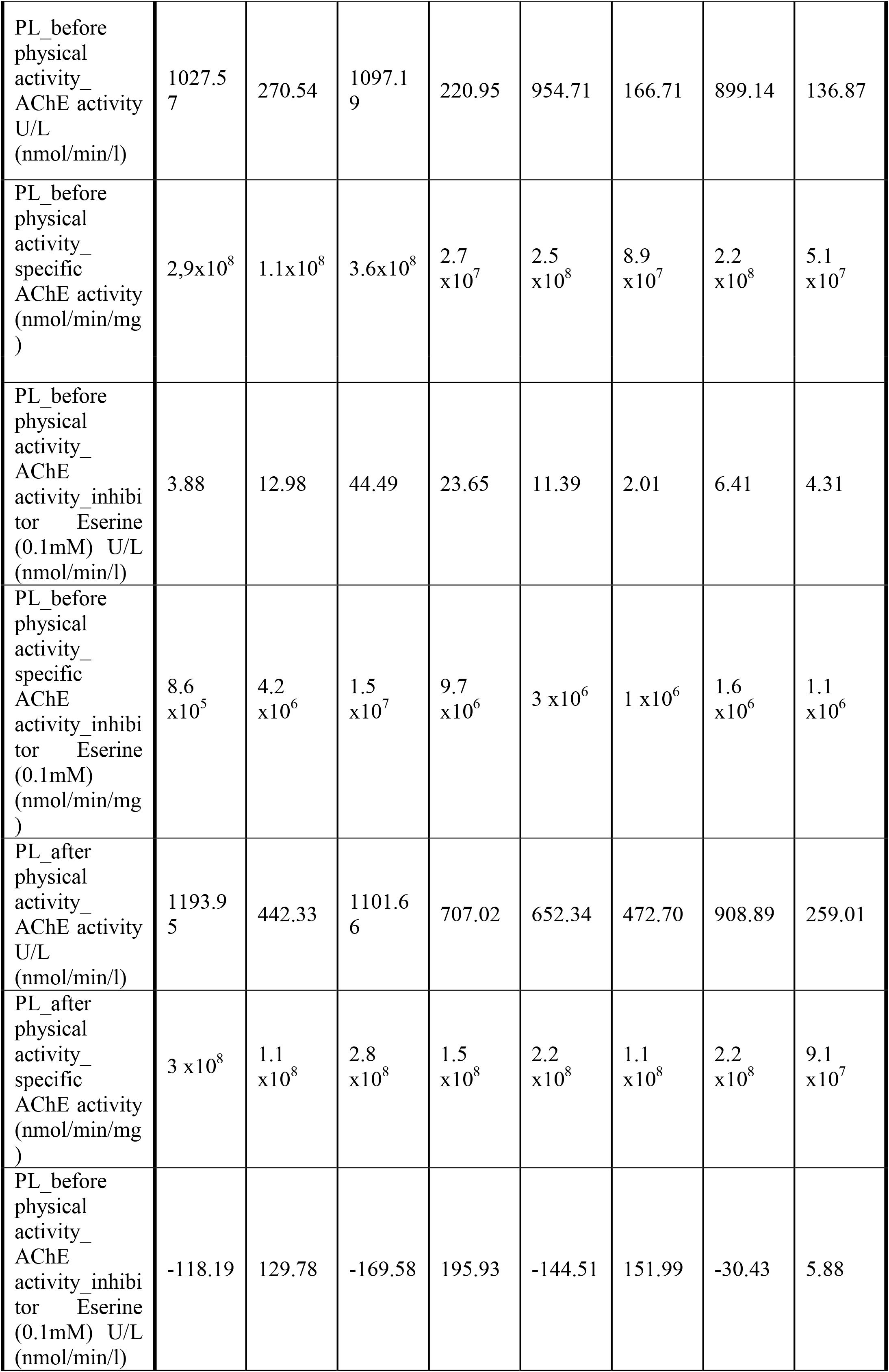
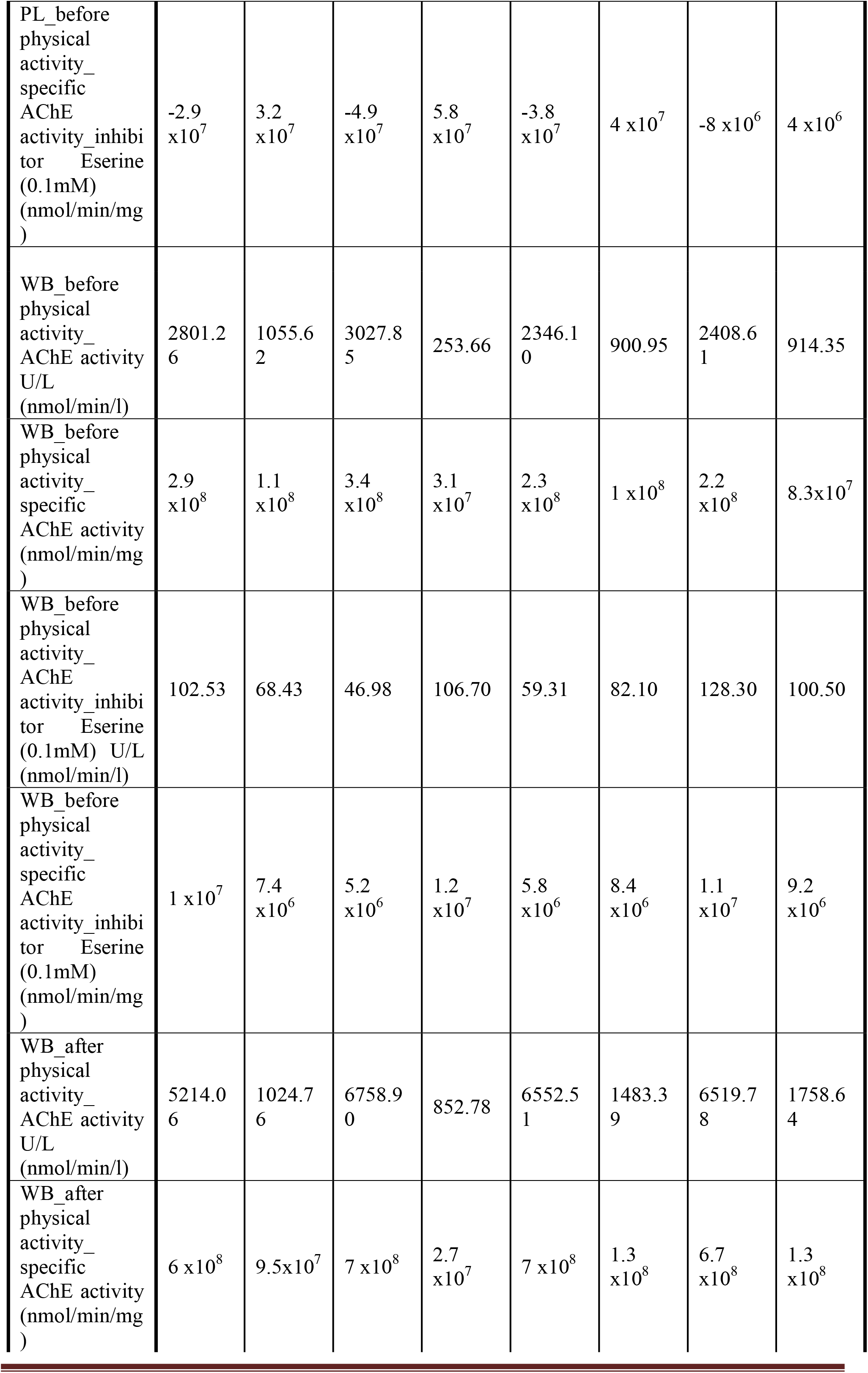
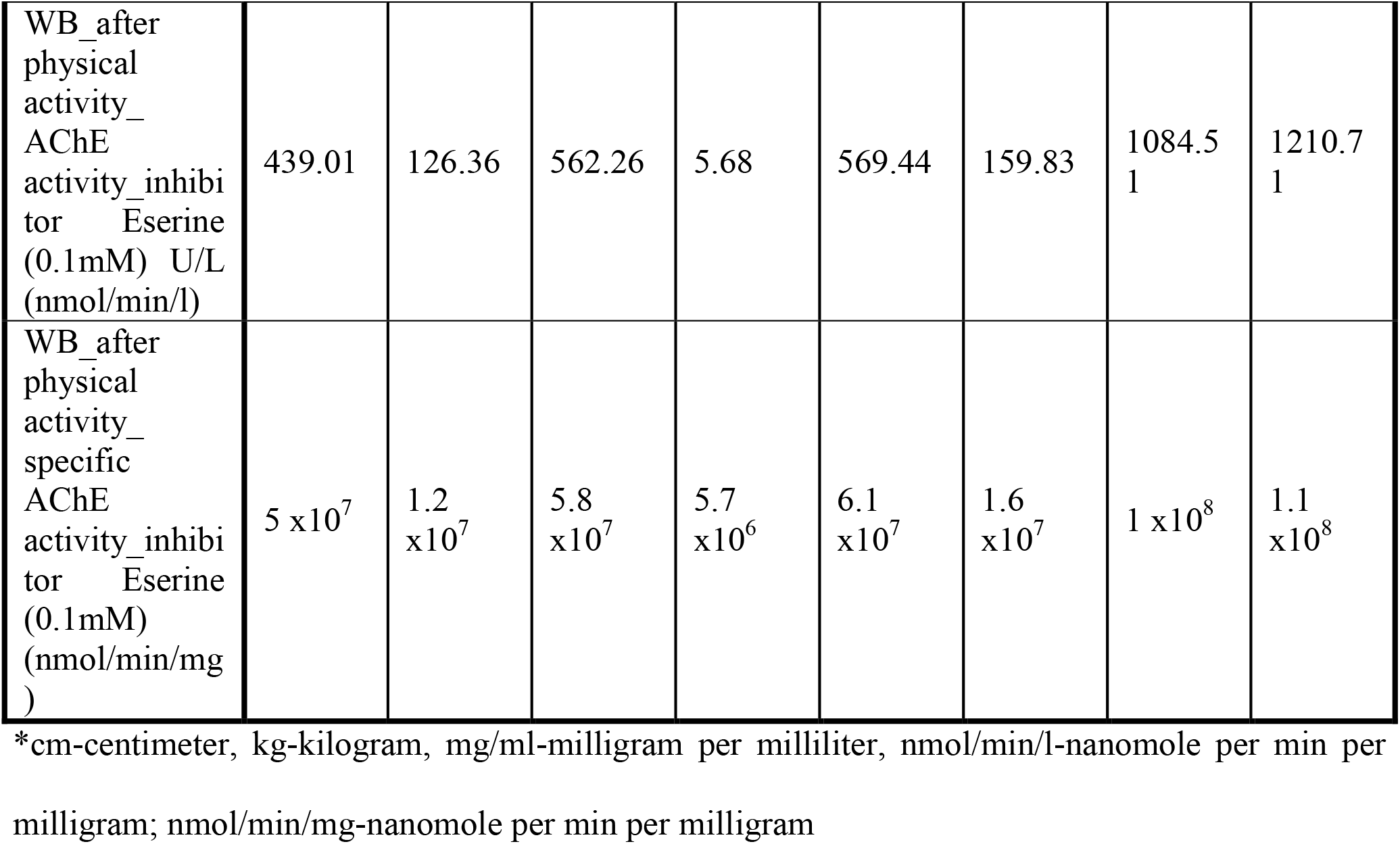
Anthropometric and Blood parameters of healthy elderly participants

**Figure 1.**
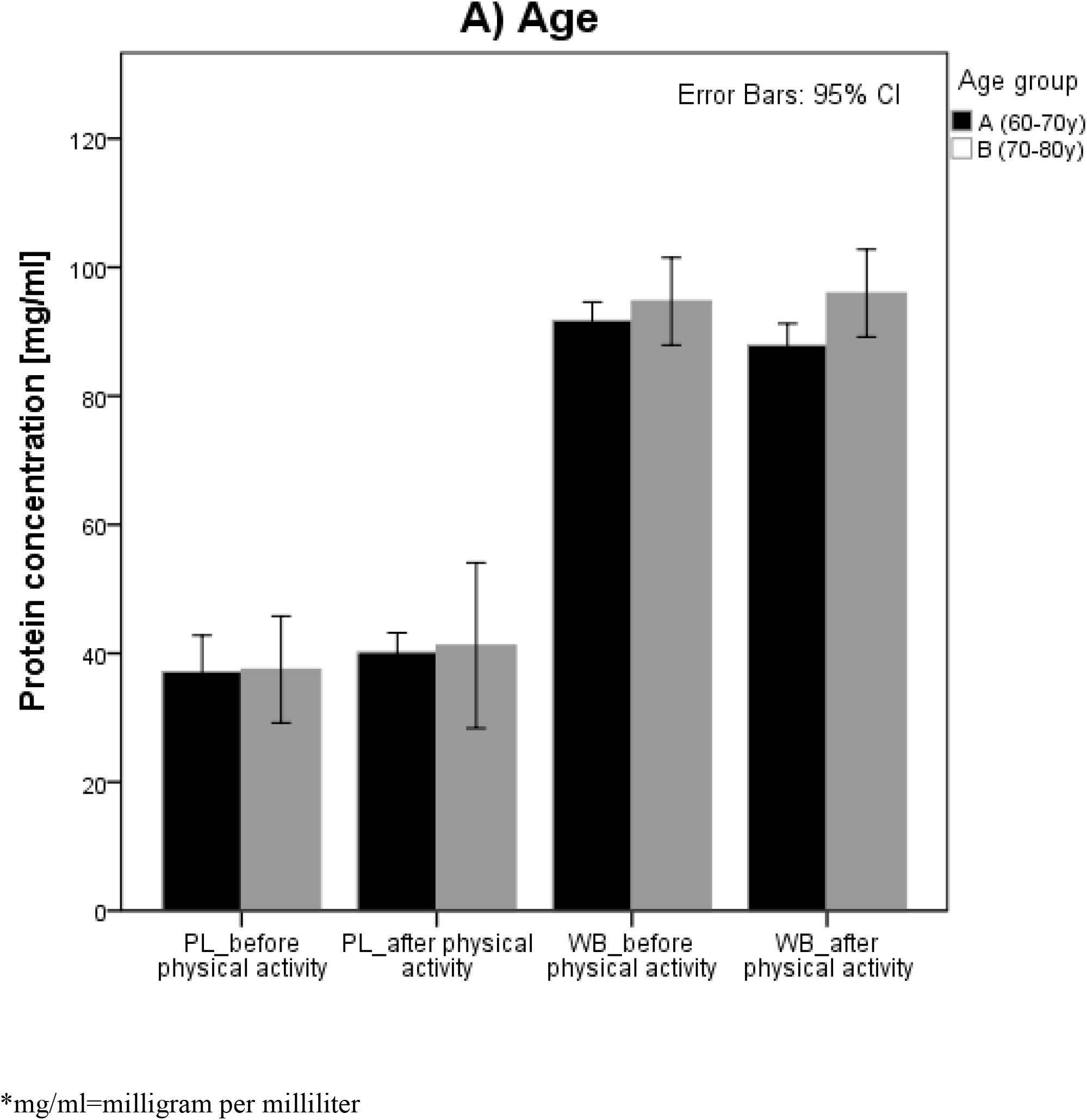
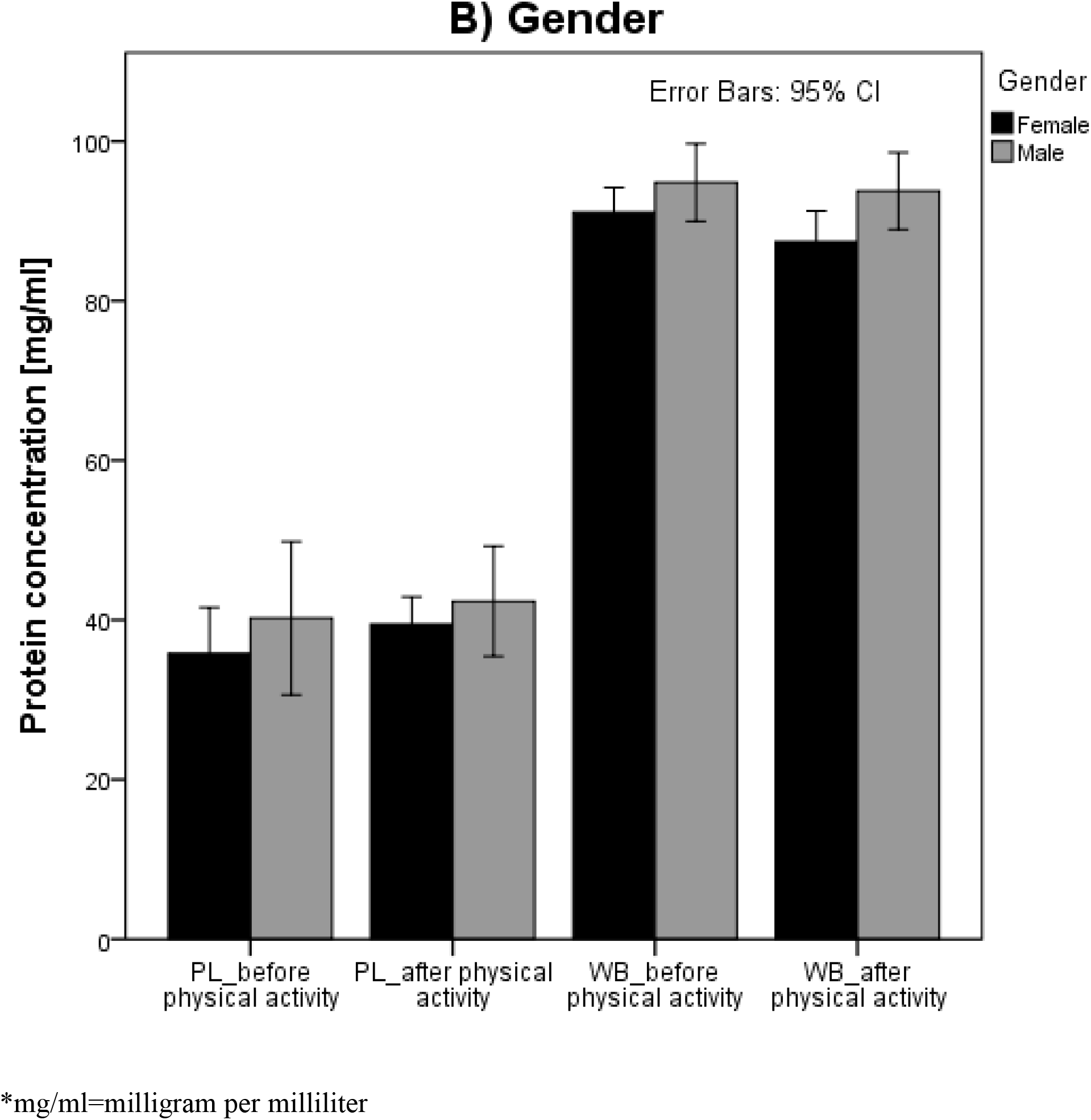
Characteristics of the healthy elderly participant’s sample group after 3 months of physical activity and storage time at −21°C. Protein concentration [mg/ml] of participants group based on Age and Gender in Plasma and Whole blood samples. Values represent investigated participants mean value of protein concentration with error bars 95% confidence interval. (mg/ml=milligram per milliliter).

**Figure 2.**
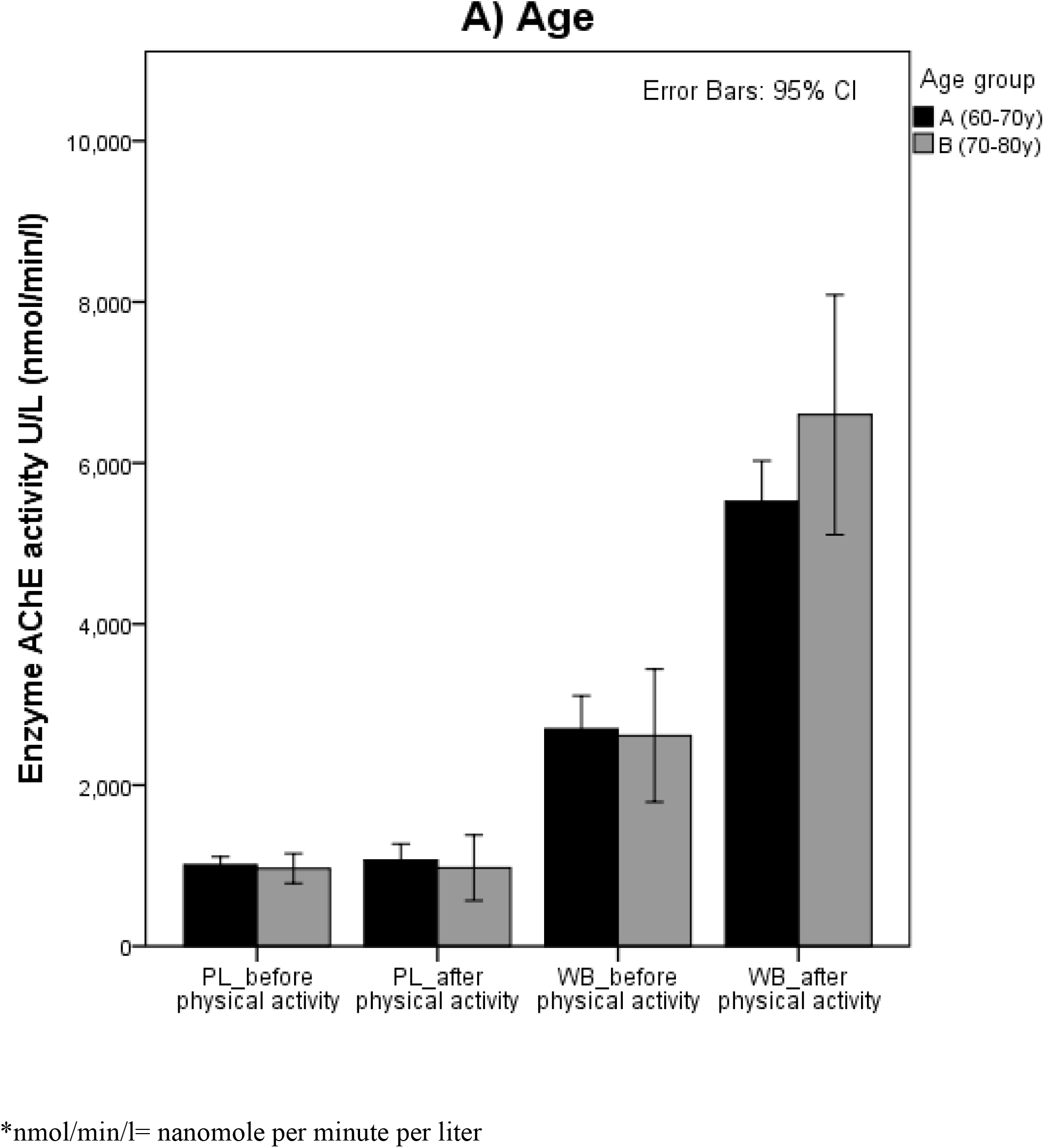
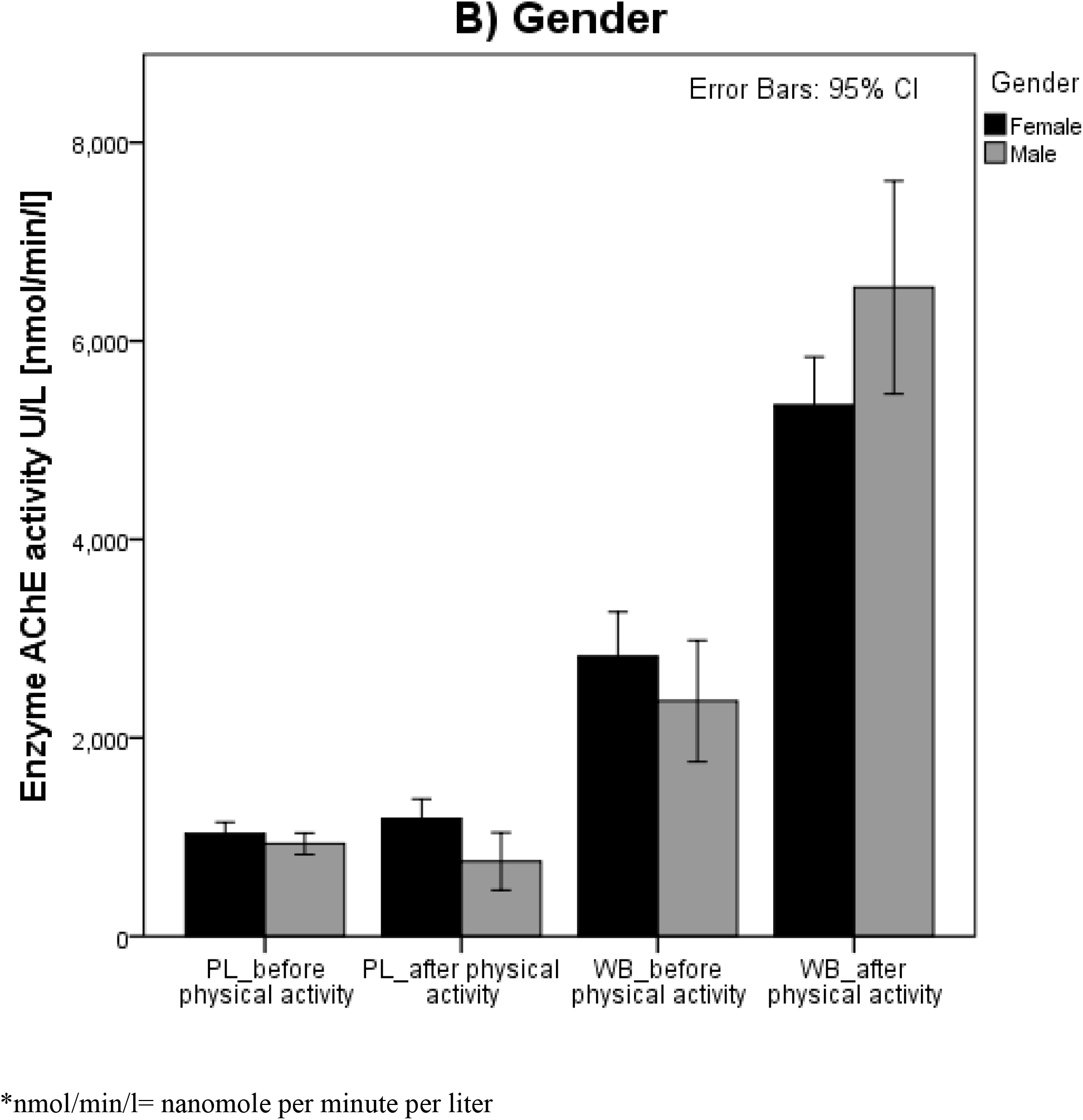
Characteristics of the healthy elderly participant’s sample group after 3 months of physical activity and storage time at −21°C. Enzyme AChE activity U/L [nmol/min/l] of participants group based on Age and Gender in Plasma and Whole blood samples using the substrate. Values represent investigated participants mean value of enzyme AChE activity with error bars 95% confidence interval. (*nmol/min/l=nanomole per minute per liter.

**Figure 3.**
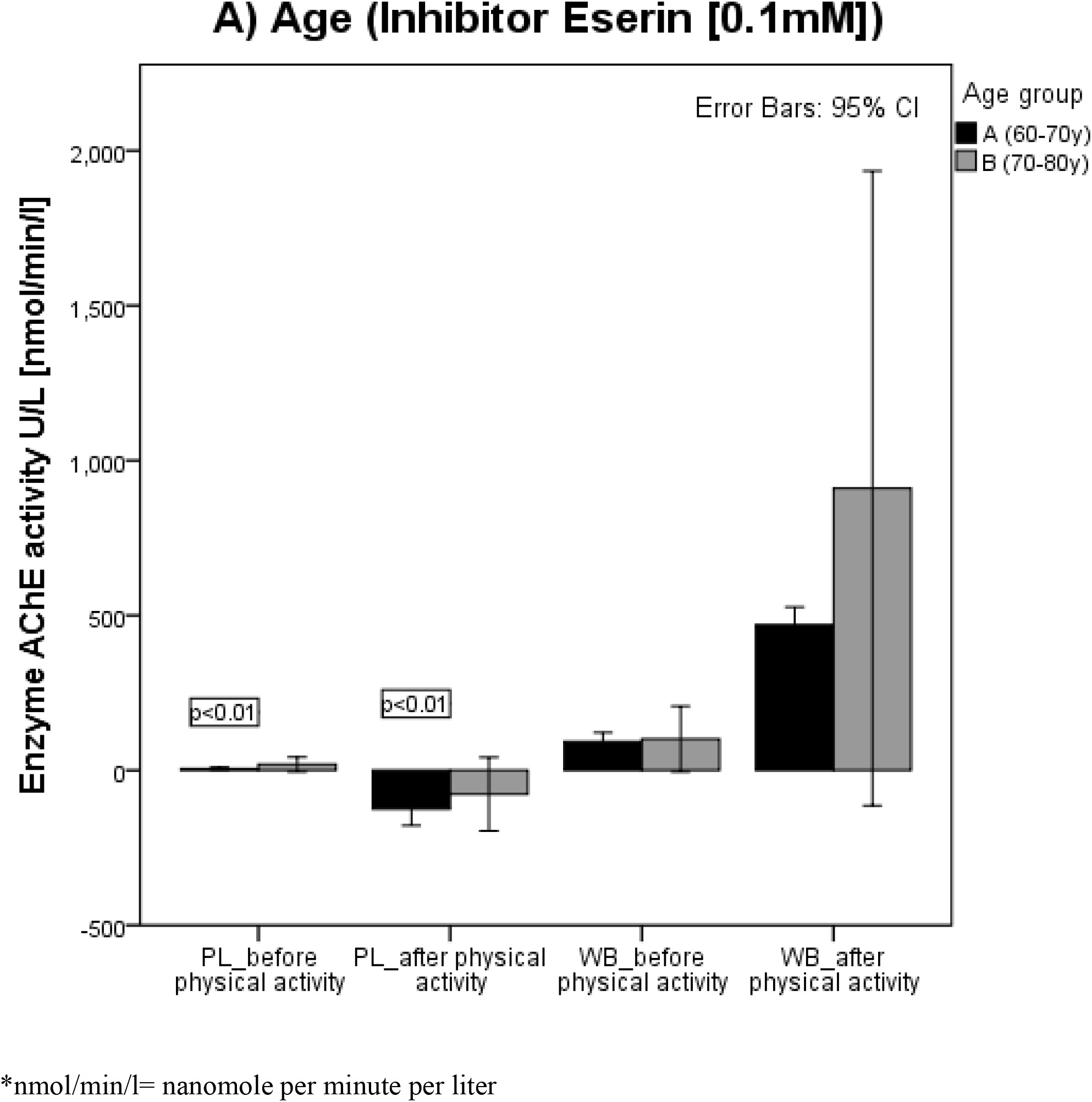
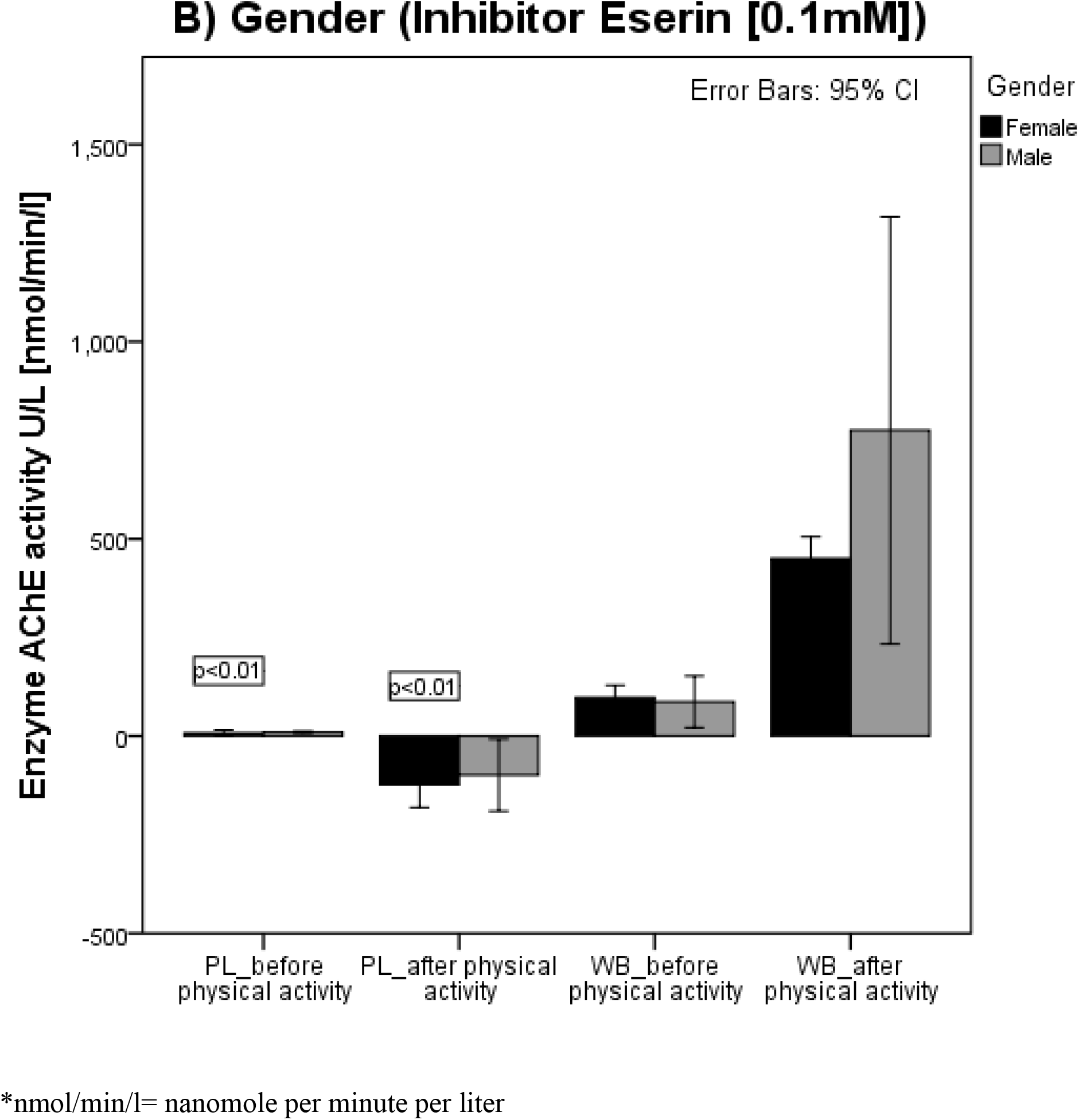
Characteristics of the healthy elderly participant’s sample group after 3 months of physical activity and storage time at −21°C. Enzyme AChE activity U/L [nmol/min/l] of participants group based on Age and Gender in Plasma and Whole blood samples using inhibitor Eserin (0.1mM). Values represent investigated participants mean value of enzyme AChE activity with error bars 95% confidence interval. (*nmol/min/l=nanomole per minute per liter).

### Assessing the significance of mean difference (Paired T-test and Wilcoxon; One sample T-test, Independent T-test and Mann Whitney/Kruskal Wallis test)

Paired T-test with a 95% confidence interval revealed a statistically significant mean difference (P<0.01) for protein concentration, basal and specific enzyme AChE activity in PL samples, inhibitor Eserine efficacy inhibition basal and specific enzyme AChE activity in WB samples before and after physical activity. Differences between PL and WB samples exist (P<0.01) for protein concentration, enzyme AChE activity and inhibitor efficacy before and after physical activity. Wilcoxon test revealed a statistically significant difference (P<0.01) for inhibitor Eserine inhibition efficacy of basal and specific enzyme AChE activity in PL samples before and after physical activity. One sample T-test (Kolmogorov-Smirnov) showed a significant mean difference for Age (P<0.01), Body Mass Index (P<0.05), Hip Circumference (P<0.05), Muscle Mass (P<0.01), inhibitor Eserine inhibition efficacy of basal and specific enzyme AChE activity in PL and WB samples before and after physical activity (P<0.01). Independent T-test showed statistical mean difference with 95% CI when grouping variable is Age (70-80 year group) for Height (P<0.01), protein concentration (P<0.05) in WB samples after physical activity. Difference between Gender is present for Height (P<0.01), Weight (P<0.05), protein concentration (P<0.05) in WB and enzyme AChE activity (P<0.05) in PL samples after physical activity. Mann Whitney and Kruskal Wallis test revealed a statistically significant difference when grouping variable is Age (70-80 year group) for Muscle mass (P<0.01) and when grouping variable is Gender for Age (P<0.05) and Muscle mass (P<0.01).

### Assessing inhibitor efficacy for inhibitor Eserine (0.1mM) and BW284c51 (0.01mM)

An experimental study, done on a total of 4 participants (3-female and 1-male), maintain the same Gender ratio from the beginning. The study assesses the inhibitor efficacy in WB and PL samples. Figure 4 (Fig.4A-C) present the difference in protein concentration, basal AChE activity and inhibitor efficacy between inhibitor Eserine and BW284c51 in PL and WB samples of healthy participants. Shapiro-Wilk test confirmed data normality for WB_protein concentration_inhibitor BW284c51 (**P**<0.05), WB_enzyme AChE activity_inhibitor Eserine (P<0.01) and WB_Eserine_Inhibitor Eserine (**P**<0.05). However, the parametric and non-parametric test showed an absence of inhibitor efficacy difference. Wilcoxon test, Unpaired T-test and Mann Whitney test showed no statistically significant mean difference between inhibitor Eserine and BW284c51 efficacies. They have the same potency. Based on this notion, we used inhibitor Eserine for all of our samples.

**Figure 4.**
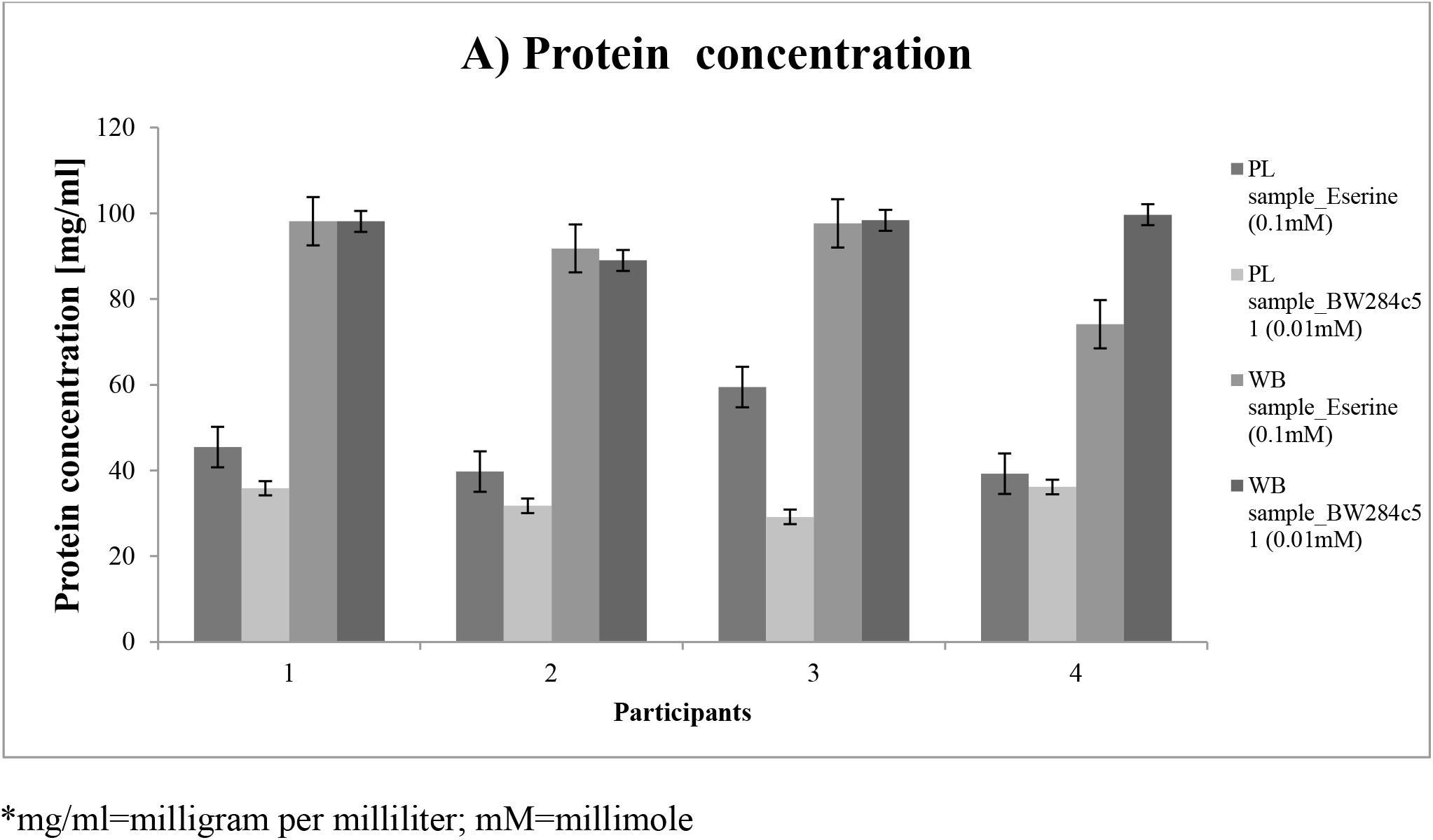
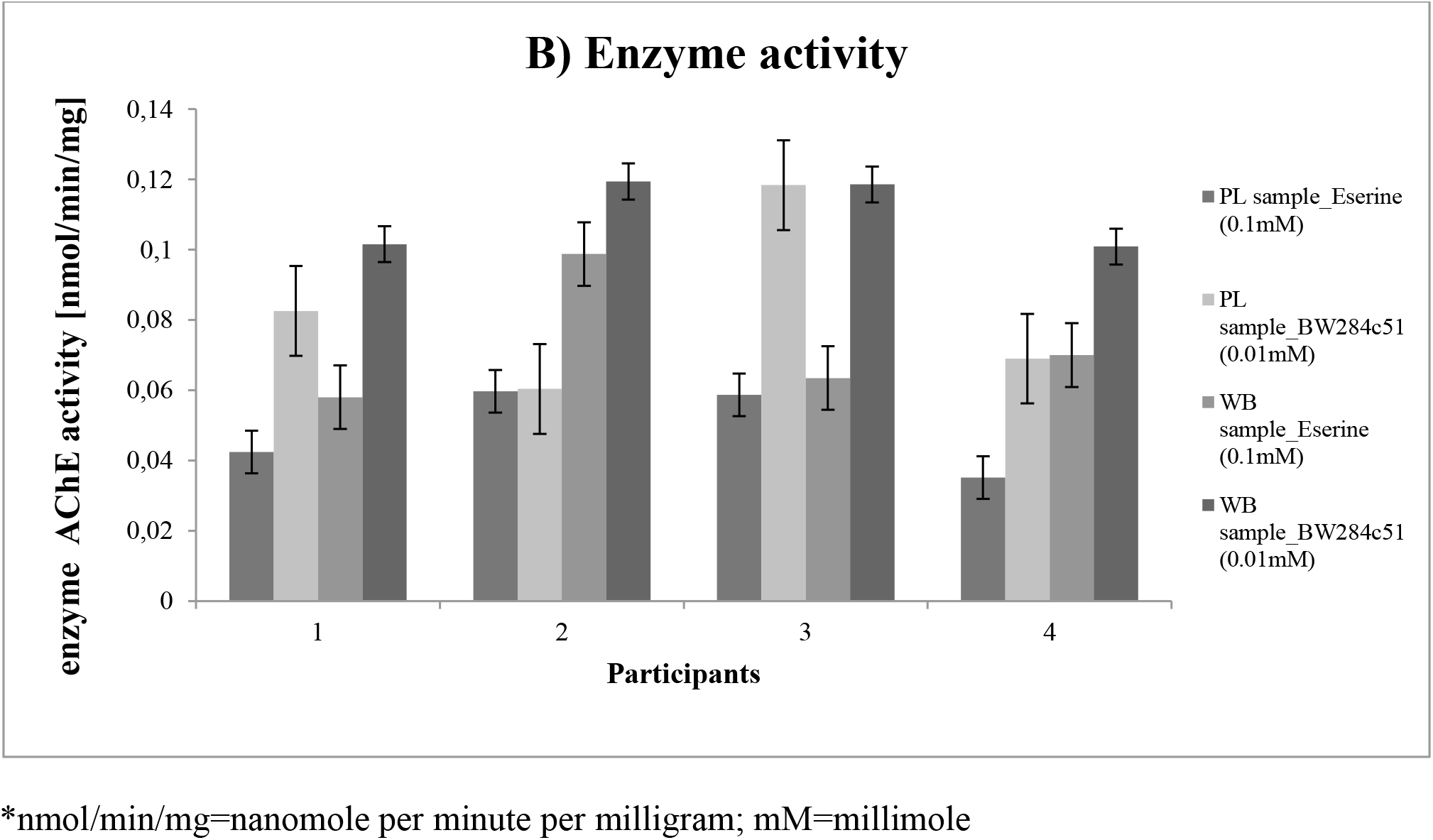
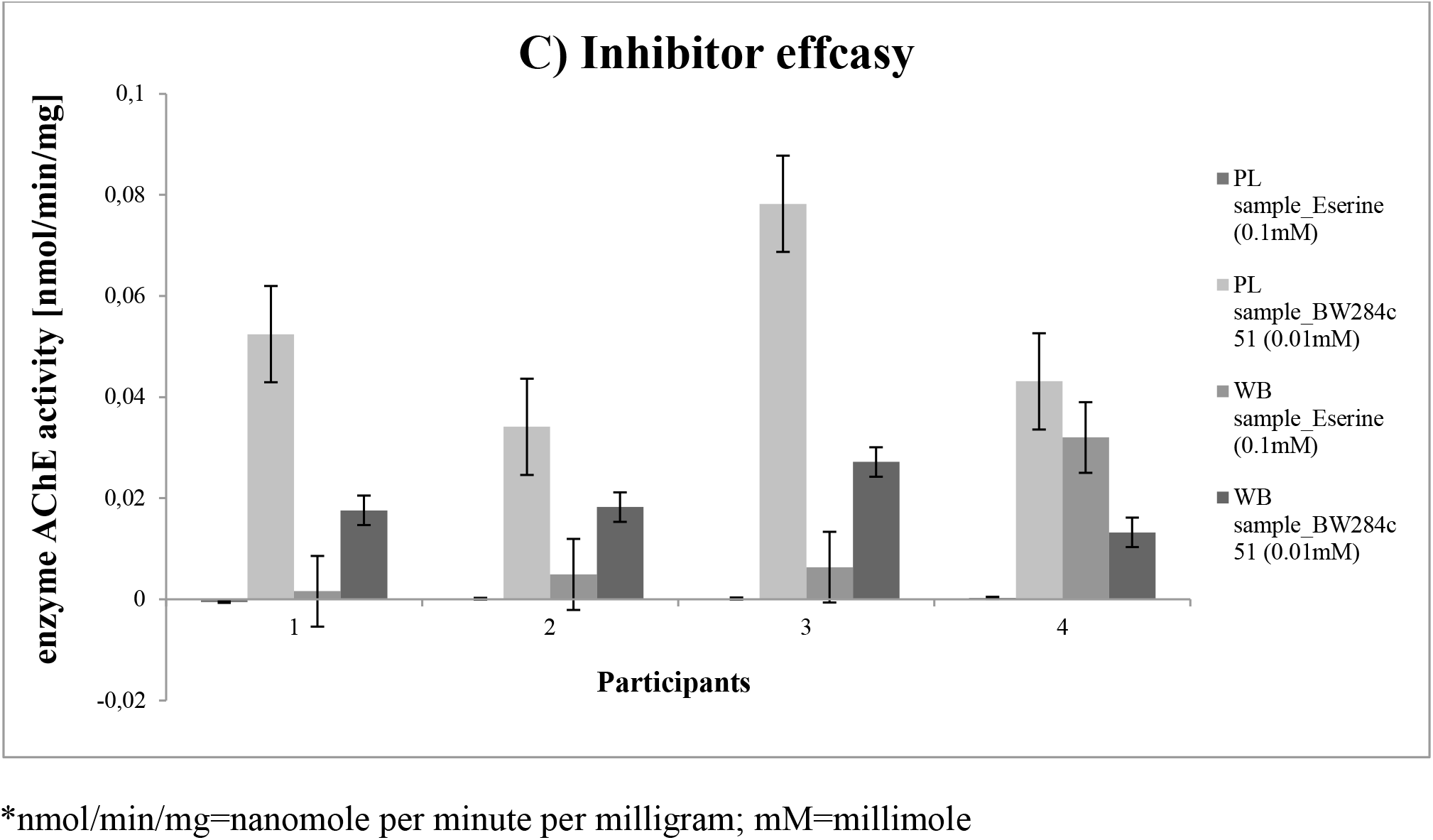
Characteristics of the healthy elderly participant’s sample group after 3 months of physical activity and storage time at −21°C. Assessing inhibitor efficacy of inhibitor Eserine (0.1mM) and BW284c51 (0.01mM) in Plasma and Whole blood samples. Values represent investigated participants mean value of protein concentration and enzyme AChE activity with error bars 95% confidence interval. (*mg/ml=milligram per milliliter; *nmol/min/l=nanomole per minute per liter).

## Discussion

Female participants had lower values in comparing to males who had higher values (% of differences) for variable Age (6.27%), Height (7.65%), Weight (15.45%) and Muscle Mass (40.85%). Results are following published data [Schorr M, 2018; Hilton EN 2021].

Information about the effect of pre-analytical factors during normal physiological conditions is essential for prevention, follow up, prognosis, treatment of disease, data interpretation and estimation of expected values during health and disease. The correlation between enzyme AChE activity during the ageing process, cancer, physical activity and storage time is known. The decline in AChE activity during normal ageing is explained by the erythrocyte membrane characteristic [Freitas Leal JK, 2017].

The level of enzyme AChE activity lacks in the literature for the elderly during physical activity. Researchers investigated the effect of 6 weeks of swimming on rats, suggesting a decrease in the enzyme AChE activity level [Cardoso AM, 2014]. Exercise improves recognition memory and AChE activity in a β-amyloid induced rat model of Alzheimer disease [Farzi MA, 2019]. In healthy human subjects, swimming increase enzyme AChE activity [Piech M, 2017]. In adult (22.5+-3.0 years) single session of physical activity does not cause the effect on enzyme AChE activity, but in adolescent (17.5+-0.6 years), it does [Zimmer KR, 2012; Schulpis KH, 2007]. Changes in enzyme AChE activity during a few days, months and years of storage is known for healthy subjects [Freitas Leal JK, 2017; Crane CR, 1970]. Discrepancies about enzyme AChE stability exist depending on the year of publishing and storage time [Muderhwa J, 2015; Gupta VK, 2015; Huizenga JR, 1985; Worek F, 1999]. AChE activity change during storage time in elderly lack in the literature for the elderly.

Current literature indicates the importance of enzyme AChE in health and disease and the absence of information regarding pre-analytical factor (monthly physical activity and sample storage freezing time) influence on protein concentration, AChE activity and inhibition inhibitor efficacy of PL and WB samples. This study compared the effect of 2 months (47 days) sample storage for samples sampled at the beginning of the physical activity (1st sampling) and 25 days of sample storage after 3. months of physical activity (2nd sampling) at −21°C in the freezer in healthy elderly individuals. Results indicate a percentage increase of enzyme AChE activity (4.9%) and a drop of specific enzyme AChE activity (0.8%) in PL samples. Inhibitor Eserine inhibits basal and specific AChE activity in PL (1517.17%/1417%) and WB (486%/528.07%) after 47 and 25 days of storage and 3. months of physical activity in healthy elderly individuals. Results conclude that 3. months of physical activity and storage time keep enzyme AChE in PL samples high (4.9%), where females have higher activity (44.38%) comparing to males, regardless of Age. Percentage of decrease for 1st and 2nd sampling after phlebotomy is not present before sample freezing. Samples of WB have higher protein concentration (85.1%/75.57%) in males aged 70-80 years and enzyme AChE activity (91.16%/137.96%) regardless of Age and Gender. Inhibitor Eserine has a higher efficacy in PL samples (168.17%/130.8%) before and after physical activity, nevertheless of Age and Gender. Slower enzyme AChE activity increases after 3. months of physical activity in the elderly group can be explained by un-reactivity of aged AChE, followed by a dealkylation reaction (ageing). Un-reactivity is a result of an evolution of the catalytic power of enzyme [Quinn DM, 2017]. The ageing process leads to irreversible AChE inhibition, resulting in the ACh accumulation of excess at the synaptic clefts [Stanciu GD, 2019].

AChE inhibitors are widely applicable in geriatric medicine for therapy [Stanciu GD, 2019; Eldufani J, 2019]. This study showed that the efficacy of inhibitor Eserine (0.1mM) and BW284c51 (0.01mM) remain the same before and after 3. months of physical activity and storage time. This finding is not following another study with inhibitor Eserine and BW284c51 [Nunes B, 2017].

Inhibitor Eserine is more efficient in WB samples (168.17%/130.84%) before and after physical activity and storage time comparing to PL samples, regardless of Age and Gender. These findings confirm the presence of a higher amount of enzyme AChE in WB than in PL. WB contains erythrocytes, leucocytes, platelets and other blood constituents rich with the enzyme AChE content.

The disadvantage of the current study is the small number of participants, Gender discrepancies and not measuring the concentration of enzyme AChE after 1st and 2nd sampling before freezing and absence of disease group. The values of enzyme AChE are not markdown before freezing. We are not able to compare the difference of enzyme AChE between 0 and 47/25 days of storage. Future analysis should include more participants. Include blood and urine parameters to compare the results on the population level. Discover additional biomarkers, specifically activated, during physical activity and affected by sample storage time. These will enable a step closer to appropriate disease treatment and therapy by tailoring a specific training regime for patients with diverse disease states. Emphasize the interaction between health and disease, including disease group and the significant influence of pre-analytical factors. However, many challenges remain to resolve since physical activity causes complex interactions with organ systems and blood, activating diverse signal network with protective and harmful mechanism.

## Conclusion

Physical activity represents a promising strategy for preventing disease onset, progression and therapy outcome in healthy elderly participants. Enable long term care and an age-friendly environment. Storage time affects enzyme activity. Physical activity has a higher impact on preserving enzyme increase, regardless of storage time. This preliminary and descriptive study shows that three months of physical activity in elderly subjects increase enzyme AChE activity in WB more than in PL samples. The study highlights the importance of the pre-analytical variables in the final result interpretation. Include simple, fast, reliable and economic methodology. Study results are applicable and repeatable for further clinical studies and early disease detection.

## List of abbreviations

ACh: acetylcholine
AChE: Acetilcholineesterase
BSA: bovine serum albumin
DTNB: The 5,5′-dithiobis (2-nitrobenzoic acid)
PL: Plasma
WB: Whole blood

## Declarations

## Acknowledgements

This manuscript has been released as a preprint at [BioRxiv platform], [Jovicic S. Association of protein concentration and enzyme AChE activity in Plasma and Whole blood samples with physical activity in healthy elderly individuals. BioRxiv 2020.09.16.299610; https://doi.org/10.1101/2020.09.16.299610. However, the current work is an upgrade of the preprint manuscript. Results of the study were presented clearly, honestly, and without fabrication, falsification, or inappropriate data manipulation.

I want to acknowledge the kind support of my CEEPUS-freemover mobility mentor from Slovenia, prof. Dr Damjana Drobne. Prof. Dr Damjana gave me the opportunity to work on this subject by ensuring me samples and place between the research rooftop. She was kind to accept me to work with her on a project, to finalize my PhD thesis through CEEPUS freemover mobility network, and establish close collaboration between Universities. She introduced me closer to academia and scientific research. I also want to acknowledge my dear colleges from the Biotechnical faculty, M.Sc Alenka Malovrh, PhD student Neža Repar, and other team members of Bionanoteam who introduced me to practical scientific work in the laboratory and student mentoring. Prof. Dr Veronika Kralj Iglic PI gave me information about the project, J5-7098, on which I worked through CEEPUS freemover mobility. I want to thank all the people who contributed to my education. My parents, for the continuously love and support. Thank you all for being a part of this lovely Snežana (Snow White) story, my life story. Life writes novels.

## Author’s information

Snežana Jovičić, a PhD student, is born on 17.10.1990 in Belgrade, Serbia. Academic qualifications: enrolment of PhD studies, Genetics (2014). Snežana Jovičić finished M.Sc. Human Molecular biology, (2014) and B.Sc. Molecular Biology and Physiology (2013).

## References

Cardoso AM, Abdalla FH, Bagatini MD, Martins CC, Fiorin Fda S, Baldissarelli J… Schetinger MR et al. (2014) Swimming Training Prevents Alterations in Acetylcholinesterase and Butyrylcholinesterase Activities in Hypertensive Rats. Am J Hypertens 27(4), 522–9. (doi: 10.1093/ajh/hpt030)

Chang EH, Chavan SS, Pavlov VA (2019). Cholinergic Control of Inflammation, Metabolic Dysfunction, and Cognitive Impairment in Obesity-Associated Disorders: Mechanisms and Novel Therapeutic Opportunities. Front Neurosci 13, 263. (doi: 10.3389/fnins.2019.00263)

Cinar D, Tas D (2015) Cancer in the elderly. North Clin Istanb 2(1), 73–80. (doi: 10.4103/apjon.apjon_52_17)

Crane CR, Sanders DC, Abbott JK (1970) Studies on the storage stability of human blood cholinesterases: I. United States Office of Aviation Medicine. AM 70-4; DOT/FAA/AM-70/4. https://rosap.ntl.bts.gov/view/dot/20893. Accessed 19 September 2020.

Eldufani J, Blaise G (2019) The role of acetylcholinesterase inhibitors such as neostigmine and rivastigmine on chronic pain and cognitive function in aging: A review of recent clinical applications. Alzheimers Dement (N Y) 5, 175–83. (doi: 10.1016/j.trci.2019.03.004)

Ellman GL, Courtney KD, Andres V, Featherstone RM. (1961) A new and rapid colorimetric determination of acetylcholinesterase activity. Biochem Pharmacol 7, 88–95. (doi: 10.1016/0006-2952(61)90145-9)

Farzi MA, Sadigh-Eteghad S, Ebrahimi K, Talebi M. (2019) Exercise Improves Recognition Memory and Acetylcholinesterase Activity in the Beta Amyloid-Induced Rat Model of Alzheimer’s Disease. Ann Neurosci 25(3), 121–5. (doi: 10.1159/000488580)

Ferlay J, Colombet M, Soerjomataram I, Dyba T, Randi G, Bettio M,…, Bray F. (2018) Cancer incidence and mortality patterns in Europe: Estimates for 40 countries and 25 major cancers in 2018. Eur J Cancer 103, 356–87. (doi: 10.1016/j.ejca.2018.07.005)

Freitas Leal JK, Adjobo-Hermans MJW, Brock R, Bosman GJCGM (2017) Acetylcholinesterase provides new insights into red blood cell ageing in vivo and in vitro. J Blood Transfus 15(3), 232–8. (doi: 10.2450/2017.0370-16)G

Gupta VK, Pal R, Siddiqi NJ, Sharma B (2015) Acetylcholinesterase from Human Erythrocytes as a Surrogate Biomarker of Lead Induced Neurotoxicity. Enzyme Res 370705. (doi: 10.1155/2015/370705.)

Hilton EN, Lundberg TR (2021) Transgender Women in the Female Category of Sport: Perspectives on Testosterone Suppression and Performance Advantage. Sports Med 51(2), 199–214. (doi: 10.1007/s40279-020-01389-3)

Huizenga JR, Van der Belt K, Gips CH (1985) The effect of storage at different temperatures on cholinesterase activity in human serum. J Clin Chem Clin Biochem 23(5), 283–5. (doi: 10.1515/cclm.1985.23.5.283)

Islam MS, Hasan MM, Wang X, Germack HD, Alam MN. (2018) A Systematic Review on Healthcare Analytics: Application and Theoretical Perspective of Data Mining. Healthcare (Basel) 6(2), 54. (doi: 10.3390/healthcare6020054)

Joyner MJ, Casey DP (2015) Regulation of increased blood flow (hyperemia) to muscles during exercise: a hierarchy of competing physiological needs. Physiol Rev 95(2), 549–601. (doi: 10.1152/physrev.00035.2013)

Lockridge O, Norgren RB Jr, Johnson R, Blake TA. (2016) Naturally Occurring Genetic Variants of Human Acetylcholinesterase and Butyrylcholinesterase and Their Potential Impact on the Risk of Toxicity from Cholinesterase Inhibitors. Chem Res Toxicol 29(9), 1381–92. (doi: 10.1021/acs.chemrestox.6b00228)

Muderhwa J, Pfluger K, Olson D, Pee A, Quintana M, Hall S. (2015) Study on the Storage Viability of Human Red Blood Cell Cholinesterase. American J Clin Path 144(2), A062. (doi: 10.1093/ajcp/144.suppl2.062)

Nunes B, Resende ST (2017) Cholinesterase characterization of two autochthonous species of Ria de Aveiro (Diopatra neapolitana and Solen marginatus) and comparison of sensitivities towards a series of common contaminants. Environ Sci Pollut Res Int 24(13), 12155–67. (doi: 10.1007/s11356-017-8761-7)

Parker M, Bucknall M, Jagger C, Wilkie R. (2020) Population-based estimates of healthy working life expectancy in England at age 50 years: analysis of data from the English Longitudinal Study of Ageing. Lancet Public Health 5, e395–403. (doi: 10.1016/s2468-2667(20)30114-6)

Piech M, Ptaszek B, Teleglow A, Marchewka J. (2017) Enzyme activity: acetylcholinesterase and glucose-6-phosphate dehydrogenase in winter swimmers. Medical Rehabilitation 21(4), 38–42. (doi: 10.5604/01.3001.0011.6828)

Quinn DM, Topczewski J, Yasapala N, Lodge A (2017) Why is Aged Acetylcholinesterase So Difficult to Reactivate? Molecules 22(9), 1464. (doi: 10.3390/molecules22091464)

Radovanović Nenadić U, Teodorović S (2020) Public Understanding, Perceptions, and Information Sources about Bioterrorism: Pilot Study from the Republic of Serbia. Health Secur 18(1), 29–35. (doi: 10.1089/hs.2019.0046)

Ramaswami R, Bayer R, Galea S (2018) Precision Medicine from a Public Health Perspective. Annu Rev Public Health 39, 153–68. (doi: 10.1146/annurev-publhealth-040617-014158)

Schorr M, Dichtel LE, Gerweck AV, Valera RD, Torriani M, Miler KK, Bradella MA. (2018) Sex differences in body composition and association with cardiometabolic risk. Biol Sex Differ 9(1), 28. (doi: 10.1186/s13293-018-0189-3)

Schulpis KH, Parthimos T, Tsakiris T, Parthimos N, Tsakiris S. (2007) An in vivo and in vitro study of erythrocyte membrane acetylcholinesterase, (Na+, K+)-ATPase and Mg2+-ATPase activities in basketball players on alpha-tocopherol supplementation. The role of L-carnitine. Clin Nutr 26(1), 63–9. (doi: 10.1016/j.clnu.2006.03.007)

Simioni C, Zauli G, Martelli AM et al. (2018) Oxidative stress: role of physical exercise and antioxidant nutraceuticals in adulthood and aging. Oncotarget 9(24), 17181–98. (doi: 10.18632/oncotarget.24729)

Stanciu GD, Luca A, Rusu RN et al. (2019) Alzheimer’s Disease Pharmacotherapy in Relation to Cholinergic System Involvement. Biomolecules 10(1), 40. (doi: 10.3390/biom10010040)

Worek F, Mast U, Kiderlen D, Diepold C, Eyer P(1999) Improved determination of acetylcholinesterase activity in human whole blood. Clin Chim Acta 288(1-2), 73–90. (doi: 10.1016/s0009-8981(99)00144-8)

World Health Organization (2021). What is the definition of health? Available from: https://www.who.int/about/who-we-are/frequently-asked-questions Accessed: 19.02.2021.

Zimmer KR, Lencina CL, Zimmer AR, Thiesen FV. (2012) Influence of physical exercise and Gender on acetylcholinesterase and butyrylcholinesterase activity in human blood samples. Int J Environ Health Res 22(3), 279–86. (doi: 10.1080/09603123.2011.634389)

